# The BMP2 prodomain promotes dimerization and cleavage of BMP6 homodimers and BMP2/6 heterodimers in vivo

**DOI:** 10.1101/2024.06.19.599755

**Authors:** Pooja Chauhan, Yongqiang Xue, Allison L. Fisher, Hyung-Seok Kim, Jodie L. Babitt, Jan L. Christian

## Abstract

Bone morphogenetic protein 2 (BMP2) and BMP6 are key regulators of systemic iron homeostasis. All BMPs are generated as inactive precursor proteins that dimerize and are cleaved to generate the bioactive ligand and inactive prodomain fragments, but nothing is known about how BMP2 or BMP6 homodimeric or heterodimeric precursor proteins are proteolytically activated. Here, we conducted in vitro cleavage assays, which revealed that BMP2 is sequentially cleaved by furin at two sites, initially at a site upstream of the mature ligand, and then at a site adjacent to the ligand domain, while BMP6 is cleaved at a single furin motif. Cleavage of both sites of BMP2 is required to generate fully active BMP2 homodimers when expressed in *Xenopus* embryos or liver endothelial cells, and fully active BMP2/6 heterodimers in *Xenopus*. We analyzed BMP activity in *Xenopus* embryos expressing chimeric proteins consisting of the BMP2 prodomain and BMP6 ligand domain, or vice versa. We show that the prodomain of BMP2 is necessary and sufficient to generate active BMP6 homodimers and BMP2/6 heterodimers, whereas the BMP6 prodomain cannot generate active BMP2 homodimers or BMP2/6 heterodimers. We examined BMP2 and BMP6 homodimeric and heterodimeric ligands generated from native and chimeric precursor proteins expressed in *Xenopus* embryos. Whereas native BMP6 is not cleaved when expressed alone, it is cleaved to generate BMP2/6 heterodimers when co-expressed with BMP2. Furthermore, BMP2-6 chimeras are cleaved to generate BMP6 homodimers. Our findings reveal an important role for the BMP2 prodomain in dimerization and proteolytic activation of BMP6.

## Introduction

Bone Morphogenetic Proteins (BMPs) are members of the TGFβ superfamily that were originally isolated as bone inducing morphogens and subsequently found to play central roles during embryogenesis and in adult homeostasis (1). Two members of this family, BMP2 and BMP6, play a pivotal role in regulating systemic iron homeostasis (2). BMP2 and BMP6 are produced by liver endothelial cells when iron levels are high (3–6). They bind to BMP receptors and the co-receptor hemojuvelin (HJV) on hepatocytes leading to phosphorylation of SMAD1/5/8 proteins that then activate transcription of the master iron regulatory hormone, hepcidin (*HAMP)*. Mice deficient in endothelial expression of either BMP2 or BMP6 exhibit hepcidin deficiency and iron overload, underscoring the critical role of these BMPs in iron homeostasis (3, 5).

BMPs are categorized into subfamilies based on sequence homology and can signal either as homodimers or as heterodimers. Class I BMPs, including BMP2 and BMP4, are capable of heterodimerizing with class II BMPs such as BMP5, BMP6, BMP7, and BMP8 (7). Heterodimers formed between class I and class II BMPs exhibit significantly higher per molecule activity compared to homodimers of either subunit, probably due to assembly of distinct receptor complexes (8, 9). Multiple studies have shown that heterodimers composed of BMP7 together with BMP2 and/or BMP4 are the endogenous ligands responsible for early patterning events in *Drosophila*, zebrafish and mouse embryos (8, 10–12). Similarly, BMP2/6 heterodimers appear to be the key endogenous regulator of hepcidin since double endothelial *Bmp2* and *Bmp6* knockout mice show the same downregulation of *Hamp* and iron overload phenotypes as do *Bmp2* or *Bmp6* single knockout mice (5, 6, 13).

The choice of whether a given BMP will form a homodimer or a heterodimer is made within the biosynthetic pathway. BMPs are initially synthesized as inactive precursor proteins that dimerize and fold in the endoplasmic reticulum (ER). They are then cleaved by members of the proprotein convertase (PC) family to generate prodomain fragments along with the active dimeric ligand (1). Although the prodomain lacks signaling activity, it plays critical roles in ligand dimerization, folding and stability (14–17). Kinetic studies have shown that furin, one of the best-characterized PCs, prefers to cleave proproteins at the carboxy-terminal side of the optimal consensus sequence -R-X-R/K-R-, although it can also cleave following the minimal sequence - R-X-X-R- (18). BMP2 and BMP4 have two highly conserved furin consensus cleavage motifs: an optimal motif adjacent to the ligand domain (the S1 site) and a minimal motif upstream of this (the S2 site). BMP4 is sequentially cleaved at the S1 and then the S2 site, with the order of cleavage being determined by the presence of optimal and minimal furin motifs (15, 16). BMP4 homodimers remain transiently associated with the prodomain following the first cleavage and are released following the second cleavage (19), whereas BMP4/7 heterodimers remain non-covalently attached to both prodomains (20). Cleavage at both sites, and formation of the transient prodomain/ligand complex are essential for formation of a functional, stable ligand in vivo (16, 21). Unlike class I BMPs, vertebrate class II BMPs have a single conserved minimal furin motif upstream of the ligand domain. Cleavage of BMP7 following this motif generates a noncovalently associated prodomain/ligand complex (22, 23). The BMP7 prodomain fragment aids in solubilizing BMP7 homodimers by shielding hydrophobic residues present in the mature domain(22). The BMP7 prodomain also binds to components of the extracellular matrix (ECM) and this holds the ligand in an inactive conformation. However, when released from the ECM, the BMP type II receptor displaces the prodomain from the ligand, enabling the ligand to signal (24–26). Unlike BMP4 and BMP7, nothing is known about the process of proteolytic activation of BMP2 or BMP6 homodimers or BMP2/6 heterodimers.

In the current studies we determine how BMP2 and BMP6 are proteolytically processed and the role of their respective prodomains in generating active BMP2 and BMP6 homodimers and BMP2/6 heterodimers. We show that BMP2 is cleaved at both furin motifs, that the order of cleavage is opposite that of BMP4, and that cleavage of both sites is essential to generate fully active homodimers and BMP2/6 heterodimers. Furthermore, our results demonstrate that BMP6 homodimers do not form in *Xenopus* embryos and reveal an important role for the BMP2 prodomain in dimerization and proteolytic activation of BMP6.

## Results

### BMP2 is sequentially cleaved at two sites, while BMP6 is cleaved at one site, within the prodomain

We used an in vitro cleavage assay to test if BMP2 is cleaved at one or both conserved furin consensus motifs within the prodomain (S1 and S2 sites, illustrated above Fig. 1A). Murine [^35^S]BMP2 was synthesized using rabbit reticulocyte lysates, incubated with recombinant furin in vitro and cleavage products were analyzed by SDS-PAGE and autoradiography at increasing time intervals. A single 28kD prodomain band appeared at 5 minutes of incubation and persisted throughout the assay (Fig. 1A, top panel, Fig. S1B). At early time points, an 18 kD ligand band was observed and this was converted into a faster migrating 14 kD species over time (Fig. 1A, bottom panel, Fig. S1B). These results suggest that BMP2 is sequentially cleaved, initially at an upstream minimal furin motif (R-X-X-R, the S2 site) and subsequently at an optimal furin motif (R-X-K/R-R, the S1 site) adjacent to the ligand domain (illustrated in Fig. 1I). This order is opposite that of BMP4, which is cleaved first at an optimal furin motif adjacent to the ligand domain, and then at an upstream minimal furin motif within the prodomain (15, 27) (Fig. S1A).

**Figure 1.**
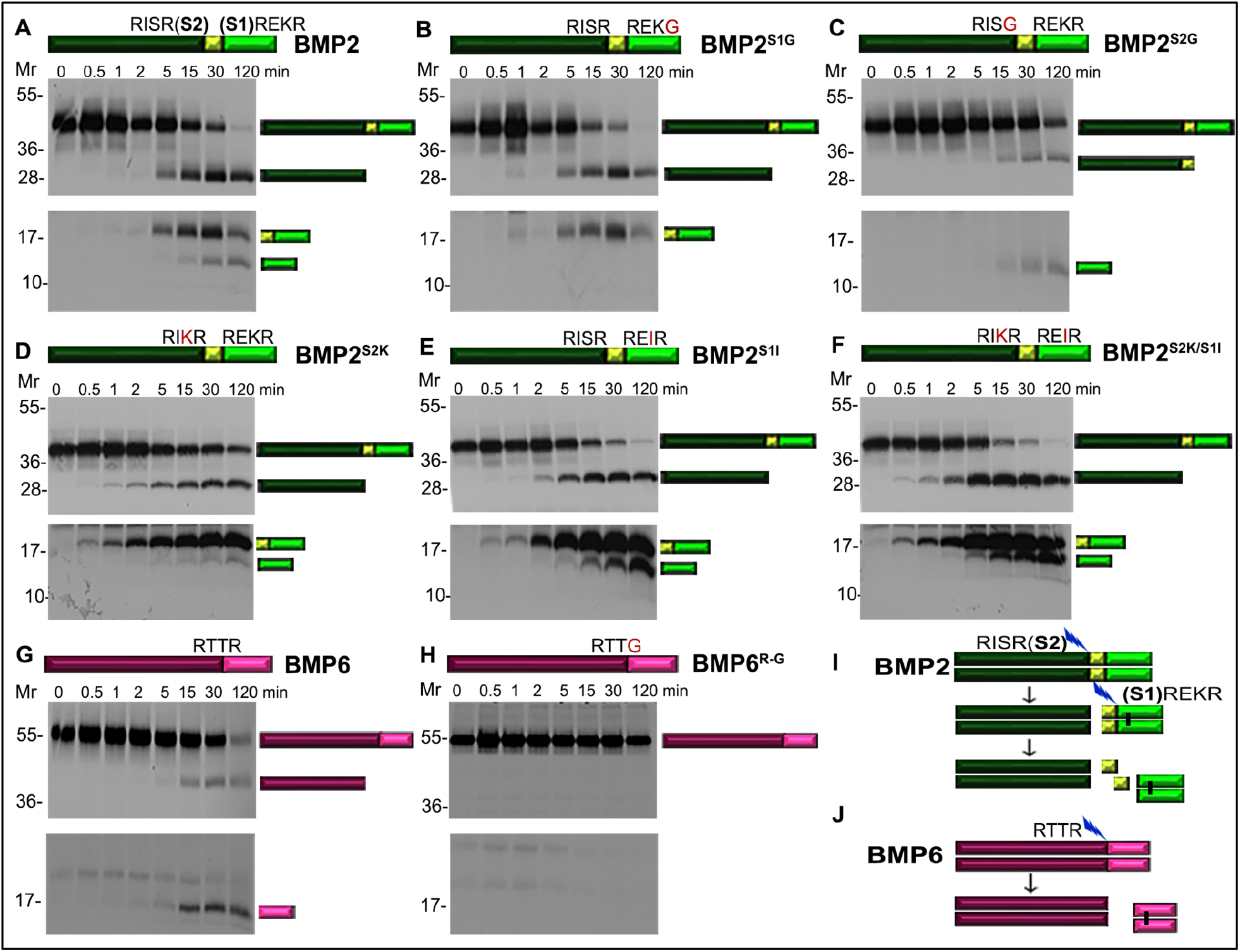
BMP2 is sequentially cleaved at two sites while BMP6 is cleaved at one site within the prodomain. (A-H) Radiolabeled precursor proteins were synthesized using rabbit reticulocyte lysate and incubated with recombinant furin. Aliquots were removed at different time points and analyzed by SDS-PAGE followed by autoradiography. Bands corresponding to the precursor, prodomain and ligand fragments are indicated to the right of the gel. In-vitro cleavage of wild-type (A) and mutant forms of BMP2 in which one of the two cleavage sites is mutated to prevent cleavage (B, C) or in which both sites are mutated to optimal (D; BMP2^S2K^) or minimal (E; BMP2^S1I^) furin motifs, or in which optimal and minimal furin motifs are switched (F; BMP2^S2K/S1I^) (illustrated above each panel. In vitro cleavage of wild-type (G) and cleavage mutant (H) BMP6 precursor protein. In A-H, top panels show a short exposure and bottom panels a longer exposure of the same gel. Fig. S1 shows a single long exposure of the intact gel shown in each panel. (I-J) Schematic illustration of cleavage of BMP2 (I), which is sequentially cleaved at the S2 and then the S1 site and BMP6 (J), which is cleaved at only one site.

To test whether cleavage of the S2 site is required for furin to recognize the S1 site, or vice versa, we analyzed mutant forms of BMP2 in which the furin consensus motif at the S1 site (BMP2^S1G^, illustrated in Fig. 1B) or the S2 site (BMP2^S2G^, illustrated in Fig. 1C) is disrupted. In vitro cleavage of BMP2^S1G^ generated a 28 kDa prodomain and an 18 kD ligand band, which appeared at the same time points and intensities as comparable bands cleaved from native BMP2 (Fig. 1B, Fig. S1C). By contrast, in vitro cleavage of BMP2^S2G^ generated faint 32 kD prodomain and 14kD ligand bands (Fig. 1C, Fig. S1D). The S2 cleaved prodomain fragment generated from BMP2 and BMP2^S1G^ first appeared at 5 minutes and both precursors were fully digested by 120 minutes (Fig. 1A, B). By contrast, the S1 cleaved prodomain fragment generated from BMP2^S2G^ first appeared at 15 minutes and high levels of uncleaved precursor persisted throughout the assay (Fig. 1C). These results suggest that furin can cleave the S2 site of BMP2 independent of, and more efficiently than, the S1 site.

Our finding that the minimal R-I-S-R motif at the S2 site of BMP2 is cleaved more efficiently than the R-E-K-R motif at the S1 site is unexpected because kinetic studies have shown that furin prefers to cleave sites that have a basic residue at the P2 position over those that do not (18). To test whether the position of minimal and optimal furin motifs influences the order of cleavage of BMP2, we analyzed in vitro cleavage of mutant forms of BMP2 containing two optimal furin motifs (BMP2^S2K^), two minimal furin motifs (BMP2^S1I^) or BMP2 in which the two sites were interchanged (BMP2^S1I/S2K^). The S2 site of all three precursor proteins was efficiently cleaved by furin and this was followed by cleavage of the S1 site. (Fig. 1D-F, Fig. S1E-G). The rate of conversion of the 18 kD ligand fragment into the 14 kD mature ligand fragment was reproducibly slower in BMP2^S2K^ (Fig. 1D, bottom panel) suggesting that the presence of an optimal motif at the S2 site hinders cleavage of the optimal motif at the S1 site. These results suggest that ordered cleavage of BMP2 at the S2 and then the S1 site is driven by sequence elements or structural features outside of the minimal and optimal furin consensus motifs.

To test whether the BMP6 precursor protein is cleaved at the single minimal furin consensus motif present in the prodomain (illustrated in Fig. 1G), we compared in vitro cleavage of native BMP6 with that of a precursor in which the consensus furin motif was disrupted (BMP6^R-G^, illustrated in Fig. 1H). Incubation of the native BMP6 precursor with furin generated a 40 kD prodomain fragment and a 17 kD ligand fragment (Fig. 1G, Fig. S1H) while the BMP6^R-G^ precursor remained intact throughout the incubation (Fig. 1H, Fig. S1I). Thus, BMP6 is cleaved following the single R-T-T-R motif to generate the active ligand (illustrated in Fig. 1J).

### Cleavage of both sites of BMP2 is required to generate fully active BMP2 homodimers and BMP2/6 heterodimers

To test if cleavage of one or both sites of BMP2 is required to generate fully active BMP2 homodimers or BMP2/6 heterodimers in vivo, we compared BMP activity in *Xenopus* embryos expressing murine BMP2, BMP2^S1G^ or BMP2^S2G^ precursor alone or together with BMP6. *Bmp* RNA was injected near the dorsal marginal zone (DMZ) of 4-cell embryos. When embryos reached the early gastrula stage the DMZ was explanted and BMP activity was analyzed by probing immunoblots of DMZ extracts with antibodies specific for pSmad1/5/8 (hereafter referred to as pSmad1, assay illustrated in Fig. 2A). We targeted ectopic BMPs to the DMZ because it has minimal endogenous Bmp signaling whereas the ventral marginal zone (VMZ) has high endogenous Bmp signaling and serves as a positive control (Fig. 2A, first two lanes of each panel). Levels of pSmad1 were significantly higher in DMZs isolated from embryos injected with RNA (25 pg) encoding BMP2 relative to those isolated from embryos injected with RNA encoding BMP2^S1G^ or BMP2^S2G^ (Fig. 2A, left panel, Fig. 2B). Levels of pSmad1 were also significantly higher in DMZs isolated from embryos co-injected with a half dose of RNA encoding BMP2 and BMP6 (12.5 pg each) relative those injected with a full dose (25 pg) of either single RNA (Fig. 2A, middle panel, Fig. 2C). This synergistic rather than additive effect when BMP2 and BMP6 are co-expressed suggests that BMP2/6 heterodimers are more active than either homodimer in *Xenopus* embryo assays as has been observed in cultured embryonic stem cells (28). Finally, levels of pSmad1 were significantly higher in DMZs isolated from embryos co-injected with RNAs encoding wild type BMP2 and BMP6 precursors relative to those isolated from embryos co-injected with RNAs encoding BMP2^S1G^ or BMP2^S2G^ together with BMP6 (Fig. 2A, right panel, Fig. 2D). Thus, cleavage of both the S1 and the S2 sites of BMP2 is required to generate fully active BMP2 homodimers and BMP2/6 heterodimers.

**Figure 2.**
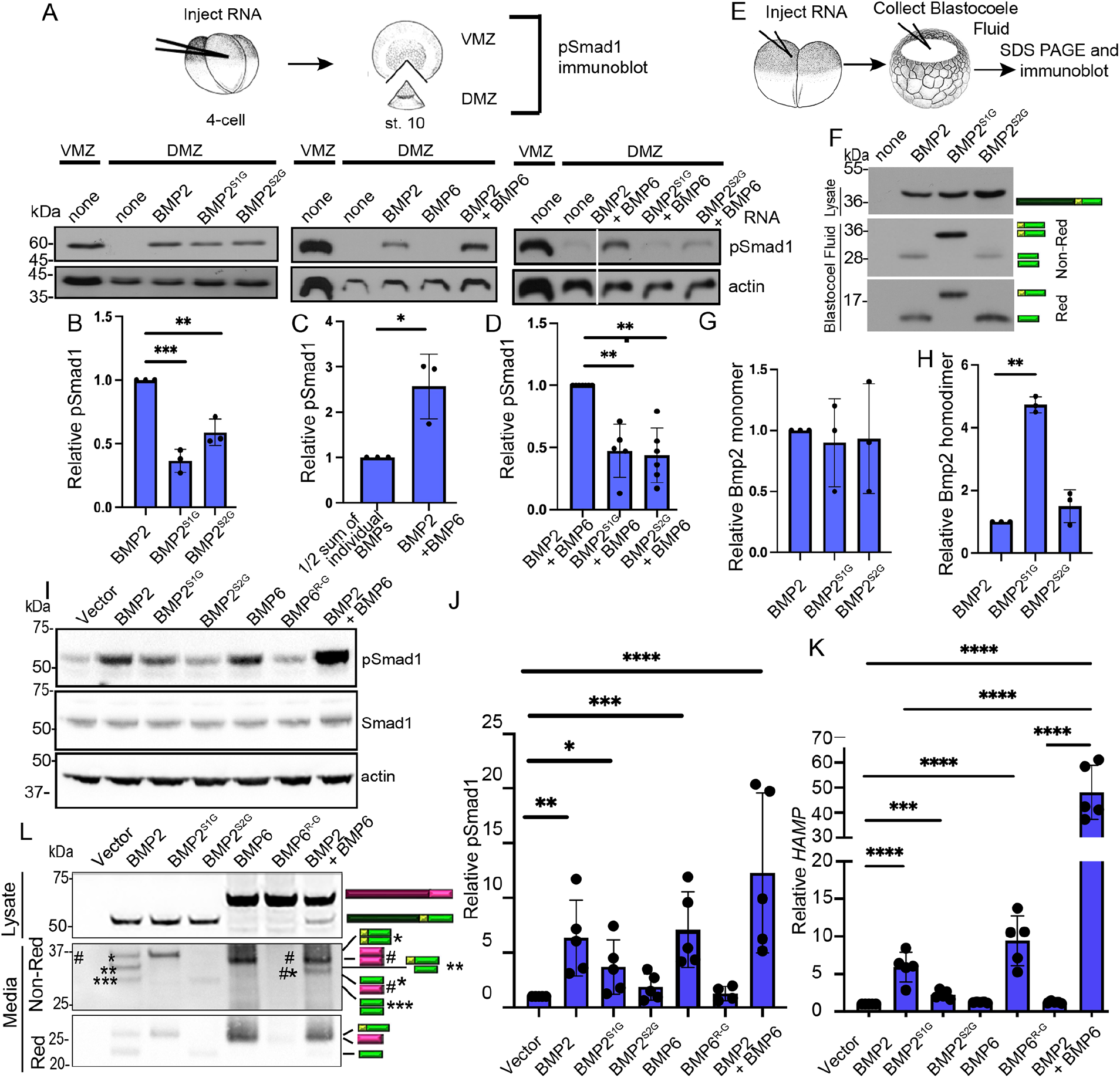
Cleavage of both sites of BMP2 is required to generate fully active BMP2 homodimers and BMP2/6 heterodimers. (A-D) RNA encoding wild-type or cleavage mutant forms of BMP2 alone or together with BMP6 was injected near the dorsal midline of four-cell *Xenopus* embryos. Ventral (VMZ) and dorsal marginal zone (DMZ) explants were collected at stage 10 and pSmad1 levels were analyzed by immunoblot as illustrated. Representative pSmad1 immunoblots in embryos expressing homodimeric (left panel) or heterodimeric (middle and right panel) ligands are shown (A). The relative level of pSmad1, normalized to actin and reported relative to that in embryos expressing wild type BMP2 alone (B), sum of individual BMP2 and BMP6 divided by 2 (to normalize RNA levels) (C) or wild type BMP2 together with BMP6 (D) is shown. Data are from three independent experiments (mean ± SD) (*p<0.05, **p<0.01, ***p<0.001 as determined by unpaired *t*-test). (E-H) Two-cell *Xenopus* embryos were injected with RNA encoding wild-type or cleavage mutant forms of BMP2. Proteins secreted into the blastocoele cavity were extracted from gastrula stage embryos and deglycosylated with PNGaseF. Immunoblots of proteins separated under reducing (red) or non-reducing (non-red) conditions were probed with antibodies specific for the HA epitope in the ligand domain as illustrated (E). Representative immunoblots are shown with bands corresponding to precursor proteins, ligand dimer and monomer indicated to the right of the gel (F). Relative levels of BMP2 monomeric (G) and dimeric (H) ligand reported relative to those in embryos expressing wild type BMP2 is shown. Data are from three independent experiments (mean ± SD) (**p<0.01 as determined by unpaired *t*-test). (I-J) TMNK1 cells were transfected with cDNAs encoding wild type or cleavage mutant BMP2 or BMP6 precursors or wild type BMP2 together with BMP6. Conditioned media was collected 24 hours later and applied to Hep3B cells for 6 hours, after which cell lysates and RNA were collected, pSMAD1 levels were analyzed by immunoblot, and *HAMP* levels were analyzed by qPCR. Representative pSMAD1 immunoblot (I). The relative level of pSMAD1 normalized to actin (J) or *HAMP* normalized to *RPL19* (K) and reported relative to that in cells transfected with empty vector is shown. Data are from a minimum of four independent experiments (mean ± SD) (*p<0.05, **p<0.01, ***p<0.001, ****p<0.0001as determined by ANOVA with Tukey post-hoc test). (L) Immunoblots of transfected TMNK1 cell lysates under reducing conditions and media under reducing (Red) or non-reducing (Non-Red) conditions probed with antibodies specific for the HA epitope in the ligand domain. Bands corresponding to precursor proteins, ligand dimers and monomers illustrated to the right of the gel.

We compared steady state levels of ectopically expressed BMP ligands in embryos expressing wild type or cleavage mutant BMP2 precursors. Two-cell embryos were injected with RNA (500 pg) encoding murine BMP2, BMP2^S1G^ or BMP2^S2G^ precursor proteins, each carrying an HA epitope tag in the ligand domain. HA-tagged BMP2 and BMP6 have the same activity as untagged proteins when ectopically expressed in mammalian HEK293T cells (Fig. S2B). At the early gastrula stage, fluid was aspirated from the blastocoel cavity, secreted proteins were deglycosylated and separated by SDS-PAGE under reducing or non-reducing conditions, and BMP2 ligands were detected by probing immunoblots with antibodies specific for the HA tag (illustrated in Fig. 2E). Equivalent levels of cleaved monomeric ligand were detected under reducing conditions in embryos expressing wild type or either of the cleavage mutant BMP2 precursors (Fig. 2F, G). Equivalent levels of BMP2 homodimers were observed in embryos expressing BMP2 or BMP2^S2G^, but higher levels of BMP2 homodimers were observed in embryos expressing BMP2^S1G^ (Fig. 2F, H). These results demonstrate that the loss of function when only one site of BMP2 is cleaved in *Xenopus* embryos is not due to lower steady state levels of ligand.

We also tested if cleavage of one or both sites of BMP2 or BMP6 is required to generate fully active homodimers in liver endothelial cells to induce expression of hepatocyte *HAMP*. TMNK1 liver endothelial cells were transfected with cDNAs encoding wild type or cleavage mutant BMP2 or BMP6 precursors. Conditioned media was collected 24 hours later and applied to the hepatocyte Hep3B cell line for 6 hours, followed by analysis of pSMAD1 levels by immunoblot and *HAMP* levels by qPCR. Conditioned media from cells transfected with BMP2 or BMP6 induced significantly higher levels of pSMAD1 and *HAMP* relative to cells transfected with empty vector (Fig. 2I-K). In contrast, only modest induction was seen in response to conditioned media from cells transfected with BMP2^S1G^, and no induction was seen for other BMP2 or BMP6 cleavage mutants (Fig. 2I-K). Immunoblot analysis of media from TMNK1 cells expressing BMP2 revealed the presence of ligand monomers corresponding to both S1 and S2 cleaved precursor under reducing conditions, and all possible cleavage intermediate dimers under non-reducing conditions with fully cleaved BMP2 dimers being the minor species (Fig. 2L, media). This contrasts with proteolytic cleavage of BMP2 in *Xenopus* embryos where only the fully cleaved BMP2 dimer is detected in blastocoel fluid at steady state (Fig. 2F). A strong band corresponding to S2 cleaved BMP2 monomer and dimer was detected in media from cells expressing BMP2^S1G^, whereas only a faint band corresponding to S1 cleaved BMP2 monomer and dimer were detected in the media of cells expressing BMP2^S2G^ (Fig. 2L, media). The latter finding is consistent with the results of in vitro cleavage showing that cleavage of the S2 site of BMP2 is required for efficient cleavage of the S1 site (Fig. 1C). Cleaved monomeric and dimeric BMP6 ligand was detected in cells expressing wild type, but not cleavage mutant BMP6 (Fig. 2L). Collectively, these results demonstrate that cleavage of BMP2 at both the S1 and the S2 site is required to generate a fully functional ligand in *Xenopus* embryos and in liver endothelial cells. Moreover, the loss of BMP activity when only S2 site is cleaved is not due to lower steady state levels of ligand, whereas inefficient S1 cleavage and lower steady state ligand levels may contribute to reduced function when the S2 site cannot be cleaved in endothelial cells.

Interestingly, despite transfection of equal amounts of cDNA in TMNK1 cells, BMP6 protein expression levels were much higher than BMP2 expression levels (Fig 2L). We therefore tested the impact of transfecting lower levels of BMP6 cDNA to try to achieve similar protein expression levels as BMP2. We found that transfection with 100-200 ng of BMP6 (1/5-1/10 dose) resulted in similar or slightly lower expression of precursor proteins and cleaved ligands under non-reducing conditions as did 1,000 ng BMP2 (Fig. S3A). Dimerized BMP2 and BMP6 precursor proteins were detected in cell media under non-reducing conditions (Fig. S3A). When similar protein levels were expressed, conditioned media from BMP6-transfected TMNK1 cells was less effective or ineffective to induce pSMAD1 and *HAMP* in Hep3B cells than conditioned media from BMP2-transfected cells (Fig S3B-D). This is consistent with the results in *Xenopus* where expression of BMP2 was less effective to induce pSmad1 than was BMP6 (Fig 2A).

Finally, we tested if BMP2/6 heterodimers can be synthesized in liver endothelial cells and signal synergistically. Media from TMNK1 cells co-transfected with one-half dose of BMP2 and BMP6 (500 ng each) induced 5 or 10-fold higher levels of *HAMP* in Hep3B cells than did media from cells transfected with full-dose BMP2 or BMP6 alone (1,000 ng each) (Fig. 2K), with similar trends seen for pSMAD1 levels (Fig. 2I-J). A synergistic effect on pSMAD1 and *HAMP* was also seen when using media from TMNK1 cells co-transfected with BMP2 and lower doses of BMP6 to achieve comparable ligand protein expression levels (Fig S3A-D). Immunoblot analysis of media from TMNK1 cells co-transfected with BMP2 and BMP6 revealed the presence of BMP6 homodimers as well as a heterodimer band that migrated at a position distinct from BMP6 homodimers or any BMP2 homodimer species (Fig. 2L, media, non-reducing).

### The prodomain of BMP2 is sufficient to generate active BMP6 homodimers, and BMP2/6 heterodimers in *Xenopus*

We next tested whether the enhanced bioactivity of BMP2/6 heterodimers relative to homodimers is intrinsic to the ligand itself, or if the prodomain of BMP2 and/or BMP6 contributes to heterodimer activity. To investigate the role of the BMP2 prodomain in ligand activity, we generated a chimeric cDNA encoding the BMP2 prodomain fused to the BMP6 mature domain (BMP2-6) (Illustrated in Fig. 3A). RNA encoding BMP2, BMP6, or BMP2-6 alone (25 pg), or a half dose of BMP2 together with a half dose of BMP2-6 or BMP6 (12.5 pg each) was injected near the dorsal midline of four-cell *Xenopus* embryos and pSmad1 levels were analyzed in DMZs isolated at the early gastrula stage. Low levels of pSmad1 were observed in DMZs isolated from embryos expressing BMP6 relative to those expressing BMP2 (Fig. 3B, left panel, C). Significantly higher pSmad1 levels were detected in DMZs isolated from embryos expressing BMP2-6 relative to those expressing the native BMP6 precursor (Fig. 3B, left panel, C). Furthermore, pSmad1 levels in DMZs isolated from embryos co-expressing BMP2-6 and BMP2 were slightly higher than those in DMZs isolated from embryos co-expressing BMP2 and BMP6 (Fig. 3B, right panel, D).

**Figure 3.**
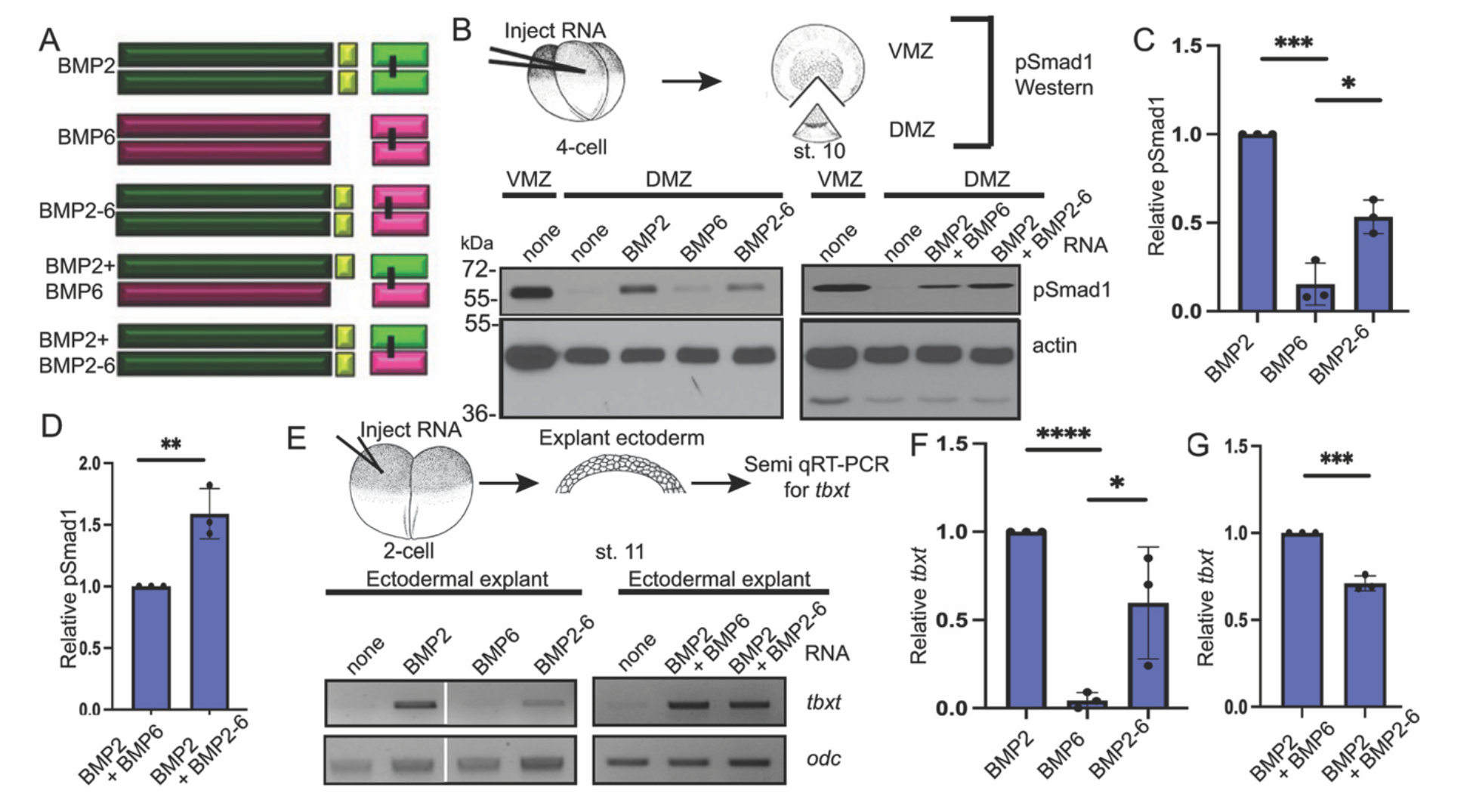
The prodomain of BMP2 is sufficient to generate active BMP6 homodimers and BMP2/6 heterodimers in *Xenopus*. (A) Schematic illustration of homodimeric and heterodimeric ligands generated from native BMP2 and BMP6 and chimeric BMP2-6 precursor proteins. (B) BMP RNAs were injected individually or together near the dorsal midline of four-cell embryos. VMZ and DMZ explants were collected at stage 10 and pSmad1 levels were analyzed by immunoblot. Representative pSmad1 immunoblots in embryos expressing homodimeric (left panel) and heterodimeric (right panel) ligands are shown. Actin levels were analyzed in duplicate samples as a loading control. (C-D) The relative level of pSmad1, normalized to actin and reported relative to that in embryos expressing native BMP2 alone (C) or together with native BMP6 (D) is shown. Data are from three independent experiments (mean ± SD) (*p<0.05, **p<0.01, ***p<0.001 as determined by unpaired *t*-test). (E) BMP RNAs were injected alone or together into one animal pole blastomere of 2-cell *Xenopus* embryos. Ectoderm was explanted at stage 11 and expression of *tbxt* was analyzed by semi qRT-PCR in embryos expressing homodimeric (left panel) or heterodimeric (right panel) ligands. All lanes from the same experiment, aligned following removal of an intervening lane (marked by white bar) using photoshop. (F-G) The relative level of *tbxt* normalized to *odc* and reported relative to that in embryos expressing native BMP2 alone (F) or together with native BMP6 (G) is shown. Data are from three independent experiments (mean ± SD) (*p<0.05, ***p<0.001, ****p<0.0001 as determined by unpaired *t*-test).

We repeated these analyses using an independent *Xenopus* ectodermal explant assay. RNA encoding BMP2, BMP6, or BMP2-6 was injected alone (50 pg), or BMP2 RNA was injected together with BMP6 or BMP2-6 RNA (25 pg each) near the animal pole of 2-cell *Xenopus* embryos. Ectoderm was explanted when the embryos reached stage 11 and levels of *tbxt* were analyzed by semi-quantitative RT-PCR (Semi qRT-PCR) (illustrated in Fig. 3E). *tbxt* is normally expressed only in mesoderm but ectopically expressed BMPs can induce *tbxt* expression in ectodermal cells (29) (Fig. 3E, left panel). Expression of *tbxt* was induced in ectodermal explants from embryos expressing BMP2 or BMP2-6, but not in explants isolated from embryos expressing BMP6 (Fig. 3E, left panel, and F). Furthermore, robust expression of *tbxt* was observed in ectodermal explants from embryos expressing BMP2 together with BMP6, or BMP2-6 together with BMP2, although the combination of BMP2 and BMP6 was slightly more potent than BMP2 and BMP2-6 in this assay (Fig. 3E, right panel, and G). These results demonstrate that the prodomain of BMP2 can enable BMP6 homodimers to signal in contexts in which the native BMP6 precursor generates little or no activity. Furthermore, BMP2/6 heterodimers generated from precursors with two copies of the BMP2 prodomain (BMP2 + BMP2-6) show similar activity to heterodimers generated from precursors containing both BMP2 and BMP6 prodomains (BMP2 + BMP6).

### The prodomain of BMP6 is not sufficient to generate active BMP2 homodimers or BMP2/6 heterodimers in *Xenopus*

To investigate the role of the BMP6 prodomain in ligand activity, a chimeric cDNA encoding the BMP6 prodomain fused to the BMP2 ligand domain (BMP6-2) was generated (illustrated in Fig. 4A). RNA encoding BMP2, BMP6 or BMP6-2 was injected alone (25 pg) or a half dose of BMP2 or BMP6-2 was injected together with a half dose of BMP6 (12.5 pg each). BMP activity was compared by analyzing pSmad1 levels in DMZs and *tbxt* levels in ectodermal explants as described previously. Significantly lower levels of pSmad1 (Fig. 4B, left panel, C) and *tbxt* (Fig. 4E, left panel, F) were detected in explants isolated from embryos expressing the BMP6-2 precursor relative to those expressing the native BMP2 precursor. Furthermore, significantly lower levels of pSmad1 and *tbxt* were observed in explants isolated from embryos co-expressing BMP6 and BMP6-2 relative to those co-expressing the native BMP6 and BMP2 precursors (Fig. 4B and E, right panels, D, G). Thus, the BMP6 prodomain is not sufficient to generate BMP2 homodimer or BMP2/6 heterodimer activity in *Xenopus*.

**Figure 4.**
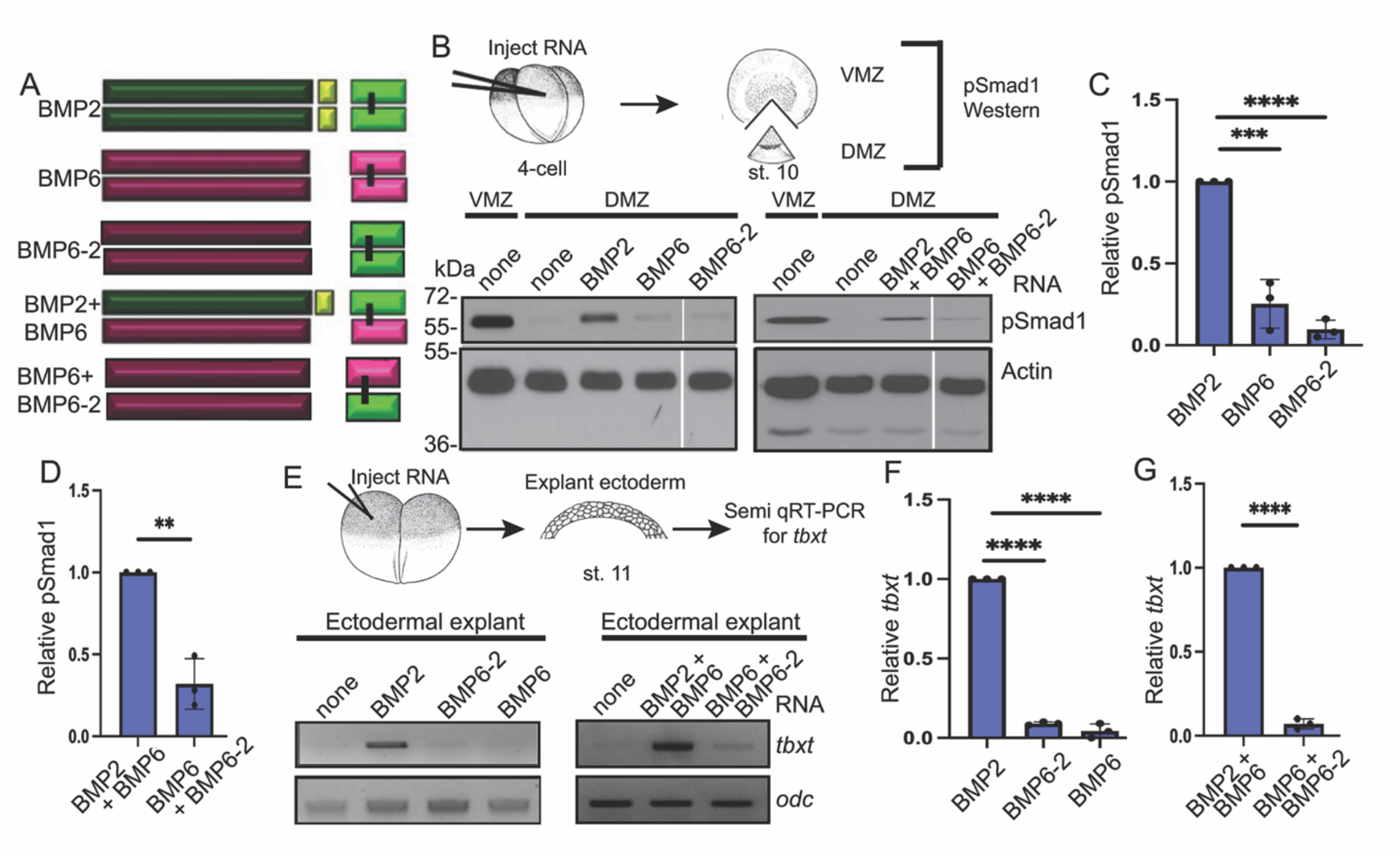
The prodomain of BMP6 is not sufficient to generate active BMP2 homodimers or BMP2/6 heterodimers in *Xenopus*. (A) Schematic illustration of homodimeric and heterodimeric ligands generated from native BMP2 and BMP6 and chimeric BMP2-6 precursor proteins. (B) RNA encoding native or chimeric BMPs were injected individually or together near the dorsal midline of four-cell embryos. DMZ and VMZ explants were collected at stage 10 and pSmad1 levels were analyzed by immunoblot. Actin levels were analyzed in duplicate samples as a loading control. Representative pSmad1 immunoblots in embryos expressing homodimeric (left panel) and heterodimeric (right panel) ligands are shown. All lanes are from the same immunoblots, aligned following removal of an intervening lane (marked by white bar) using photoshop. The pSmad1 and actin immunoblots shown in Fig. 3B and 4B are from a single injection replicate and all samples were run on the same gel. Thus, only the last lanes (BMP2-6, or BMP2 + BMP2-6 for Fig. 3B, BMP6-2, or BMP6 + BMP6-2 for Fig. 4B) differ between the panels (original blots shown in Fig. S4A). (C-D) The relative level of pSmad1, normalized to actin and reported relative to that in embryos expressing native BMP2 alone (C) or together with native BMP6 (D) is shown. Data are from three independent experiments (mean ± SD) (**p<0.01, ***p<0.001****p<0.0001, as determined by unpaired *t*-test). (E) BMP RNAs were injected alone or together into one animal pole blastomere of 2-cell embryos. Ectoderm was explanted at stage 11 and expression of *tbxt* was analyzed by semi qRT-PCR in embryos expressing homodimeric (left panel) or heterodimeric (right panel) ligands. The *tbxt* and *odc* gels shown in the left panels of Fig. 4E and 5E are from a single injection replicate and samples were run on the same gel. Thus, only the BMP2-6 or BMP2 + BMP2-6 (Fig. 3E) and BMP6-2 or BMP6 + BMP6-2 (Fig. 4E) lanes differ between the two (original gels shown in Fig. S4B). (F-G) The relative level of *tbxt* normalized to *odc* and reported relative to that in embryos expressing native BMP2 alone (F) or together with native BMP6 (G) is shown. Data are from three independent experiments (mean ± SD) (****p<0.0001 as determined by unpaired *t*-test).

### The BMP2 prodomain facilitates dimerization and cleavage of the BMP6 precursor protein in *Xenopus*

We used immunoblot analysis to examine BMP2 and BMP6 homodimeric and heterodimeric ligands generated from native and chimeric precursor proteins in *Xenopus* embryos. RNAs encoding HA-epitope tagged BMPs were injected into *Xenopus* embryos and immunoblots of proteins secreted into the blastocoele cavity were probed with antibodies specific for the HA epitope as described previously. In embryos expressing BMP2 alone, cleaved BMP2 homodimers that migrate at the predicted relative mobility (Mr) of 28 kDa were detected in blastocoele fluid under non-reducing conditions (Fig. 5A-C). On long exposures, dimerized BMP2 precursor protein was also detected in the blastocoele under non-reducing conditions (Fig. 5B). By contrast, neither the BMP6 precursor protein nor cleaved BMP6 ligand was detected in the blastocoele of embryos expressing BMP6 alone (Fig. 5A-C). In embryos expressing the BMP2-6 chimera, a band that migrates at the predicted Mr of intact dimeric BMP2-6 precursor protein (100 kDa), as well as cleaved BMP6 homodimers that migrate at the predicted Mr of 34 kDa were observed under non-reducing conditions, whereas neither precursor protein nor cleavage products were detected in the blastocoele of embryos expressing BMP6-2 (Fig. 5A).

**Figure 5.**
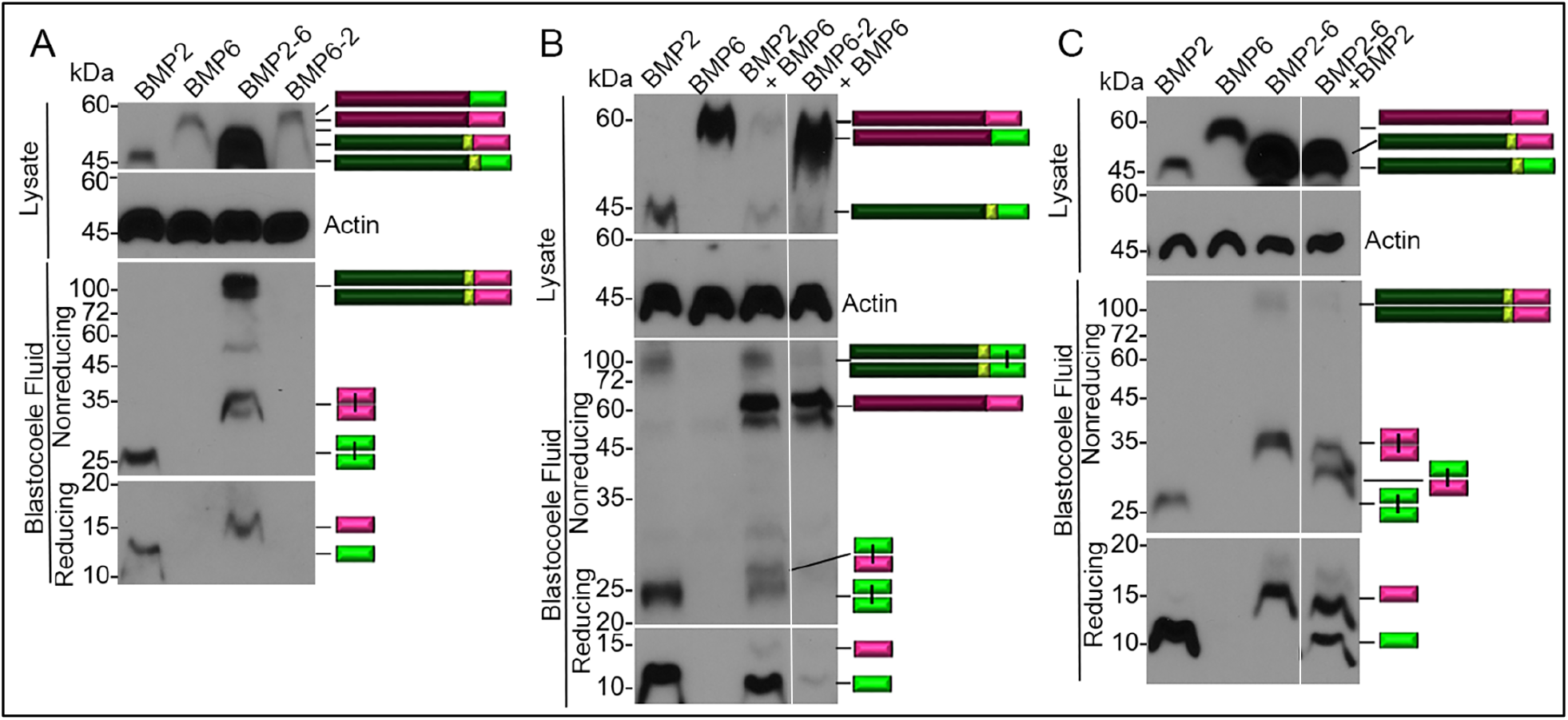
The BMP2 prodomain facilitates dimerization and cleavage of the BMP6 precursor protein in *Xenopus*. (A-C) *Xenopus* embryos were injected with RNA encoding native or chimeric forms of BMPs alone to generate homodimers (A) or in the indicated combinations to generate heterodimers (B, C). Proteins secreted into the blastocoele cavity were extracted from gastrula stage embryos and deglycosylated with PNGaseF. Immunoblots of proteins separated under reducing or non-reducing conditions were probed with antibodies specific for the HA epitope in the ligand domain. Representative immunoblots are shown with bands corresponding to precursor protein or ligand monomers or dimers indicated to the right of each gel. All lanes are from the same experiment, aligned following removal of an intervening lane (marked by white bar in B and C) using photoshop. Results were reproduced in at least three independent experiments.

When BMP2 and BMP6 precursor proteins were co-expressed in *Xenopus* embryos, cleaved BMP2 homodimers, as well as BMP2/6 heterodimers that migrate at a position intermediate between that of either homodimer were detected in the blastocoele fluid under non-reducing conditions (Fig. 5B). Monomeric BMP6 precursor protein was also detected in the blastocoele of embryos co-expressing BMP2 and BMP6 (Fig. 5B). In embryos co-expressing BMP6-2 together with BMP6, monomeric BMP6 precursor protein was secreted into the blastocoele, but cleavage products were not detected. When BMP2-6 and BMP2 were co-expressed, cleaved BMP2/6 heterodimers and BMP6 homodimers were generated (Fig. 5C). For reasons that are not clear, the BMP2-6 precursor protein is expressed at much higher levels than any other precursor protein following injection of equivalent amounts of each RNA (Fig. A, C). This may explain why co-expression of BMP2-6 and BMP2 generates heterodimers along with BMP6 homodimers while co-expression of BMP6 and BMP2 generates heterodimers along with BMP2 homodimers (Fig. B, C). Collectively, these findings support a model in which the BMP6 precursor protein cannot homodimerize or exit the ER and is not cleaved by PCs in *Xenopus* embryos. However, it appears that the BMP2 ligand domain can function in trans to facilitate secretion of the BMP6 precursor protein (BMP6-2 + BMP6; Fig. 5B). Furthermore, our results show that the BMP2 prodomain can function either in cis (BMP2-6) or in trans (BMP2 + BMP6) to the BMP6 ligand domain to facilitate dimerization, secretion and cleavage. These findings explain the absence of BMP activity in embryos expressing BMP6 or BMP6-2 chimeric precursors (Fig. 3, 4).

## Discussion

Previous studies have shown that the process of proteolytic activation and the role of the prodomain of the class I BMP, BMP4, is distinctly different than that of the class II BMP, BMP7, but it was unknown whether these processes were conserved with other members of the class I or class II BMP subfamily. Furthermore, BMP4 and BMP7 were shown to preferentially form heterodimers rather than either homodimer when co-expressed in *Xenopus* embryos, and the BMP4 prodomain was shown to be necessary and sufficient for generation of active BMP4/7 heterodimers (20). However, an unanswered question was whether all classI/II heterodimers followed a similar pattern. The current studies demonstrate that dimerization and proteolytic activation of BMP2, BMP6 and BMP2/6 precursor proteins differs considerably from that of BMP4, BMP7 and BMP4/7.

Both BMP6 and BMP7 have a single conserved minimal furin consensus motif that is recognized and cleaved by furin in vitro. However, in *Xenopus* embryos and in cultured mammalian cells the BMP7 precursor protein dimerizes and is cleaved. Furthermore, the intact BMP7 precursor is not secreted from cells unless the cleavage site is mutated to prevent recognition by PCs (20, 30). By contrast, the BMP6 precursor protein is not secreted or cleaved when expressed alone in *Xenopus* embryos. One question this raises is whether BMP6 and BMP7 are differentially trafficked through the secretory system and/or cleaved in different subcellular compartments.

BMP2 and BMP4 both have conserved minimal and then optimal furin motifs upstream of the ligand domain. For BMP4, the sequence of the motif dictates the order of cleavage since BMP4 precursor is cleaved first at the optimal motif followed by the minimal motif, whereas mutant BMP4 precursor with two optimal motifs is simultaneously cleaved at both sites (15, 19). However, despite the high degree of homology within and surrounding the cleavage sites, BMP2 is cleaved in the opposite order. The *Drosophila* ortholog of BMP2 and BMP4, Dpp, is also cleaved at multiple furin motifs. Like BMP2, the upstream site of Dpp is cleaved first, and this is essential for efficient cleavage of downstream sites (31, 32). Dpp activity is regulated by tissue-specific cleavage of the upstream site in the wing disc but not in the gut (32), raising the possibility that tissue-specific cleavage of BMP2 may provide a mechanism to differentially regulate ligand activity in vertebrates.

In some cases, class II BMP precursor proteins must heterodimerize with class I BMPs to fold and/or exit the ER, and these processes are guided by the prodomain of the class I BMP partner. In the *Drosophila* wing disc, the fly ortholog of BMP5-8, Gbb, remains in the ER in cells that express Gbb alone, but is secreted as a heterodimer from cells that co-express the BMP2 and BMP4 ortholog, Dpp (10). Vertebrate BMP4 and BMP7 preferentially form heterodimers rather than either homodimer when co-expressed in *Xenopus* embryos, and the BMP4 prodomain is both necessary and sufficient to form properly folded, functional BMP4/7 heterodimers (20). A chimeric construct consisting of the BMP4 prodomain and BMP7 ligand domain (BMP4-7) forms properly folded, functional heterodimers when co-expressed with BMP4, while the analogous BMP7-4 construct forms misfolded, inactive heterodimers when co-expressed with BMP7 (20). These findings suggest that intrinsic properties of the prodomain of Class I BMPs contribute to the bioactivity of BMP4/7 heterodimers. The current findings demonstrate that the prodomain of BMP2 also plays an essential role in formation of BMP2/6 heterodimers by facilitating dimerization and/or cleavage of the BMP6 precursor. An unanswered question is whether these roles are intrinsic to the BMP2 prodomain, or whether the BMP4 prodomain can substitute in promoting dimerization and cleavage of BMP6. Our finding that the BMP2 prodomain can function either in cis- or in trans-to BMP6 to promote dimerization and cleavage raise the question of whether the BMP2 prodomain physically interacts with the BMP6 ligand domain prior to and/or following cleavage.

The current findings demonstrate both cell type similarities and differences in proteolytic processing of BMP2, BMP6 and BMP2/6. In both *Xenopus* embryos and in mammalian liver endothelial cells, cleavage of both sites of BMP2 is essential to generate fully active homodimers. However, whereas only fully cleaved BMP2 ligand is observed in *Xenopus* embryos, ligand cleaved at the S1 or S2 site alone is observed in cultured liver endothelial cells expressing the wild type BMP2 precursor protein. Additionally, whereas the BMP6 precursor protein does not form homodimers, nor is it cleaved when expressed in *Xenopus* embryos, bioactive mature BMP6 homodimers are secreted from cultured liver endothelial cells. Whether these differences are caused by cell type specific differences in PC expression and or differences in subcellular trafficking of substrates are questions for future studies. Notably, when co-expressed in both *Xenopus* embryos and cultured liver endothelial cells, BMP2 and BMP6 form heterodimers, which exhibit increased signaling activity compared with homodimers. Although the BMP2 prodomain is necessary to promote dimerization and/or cleavage of the BMP6 precursor in *Xenopus* embryos, the BMP6 prodomain is sufficient to support dimerization and cleavage of BMP6 precursor in cultured liver endothelial cells. These data suggest that additional factors contribute to the preferential formation of BMP2/6 heterodimers in some cellular contexts, which is an important area for future investigation.

### Experimental Procedures

#### *Xenopus* Embryo Culture and Manipulation

Animal procedures followed protocols approved by the University of Utah Institutional Care and Use Committee. *Xenopus* embryos were obtained, microinjected with synthetic capped RNA and cultured as described (33). Embryo explants were performed as described (20, 34). Embryonic stages are according to Nieukoop and Faber (35).

#### cDNA Constructs

The mouse BMP6 cDNA was purchased from genscript and the mouse BMP2 cDNA was a gift from Daniel Constam. Sequence encoding an HA epitope tag was introduced into the BMP6 cDNA 23 amino acids downstream of the furin cleavage site (-V-S-R-G-[YPYDVPDYA]-S-G-S-) or into the BMP2 cDNA 6 amino acids downstream of the S1 cleavage site (-K-H-K-Q-[YPYDVPDYA]-R-K-R-L-) using the PCR-based splicing by overlap extension method (36). The same method was used to generate BMP2-6 and BMP6-2 chimeric cDNAs. cDNAs encoding point mutant forms of BMP2 and BMP6 were generated using a QuikChange II XL site-directed mutagenesis kit (Agilent Technologies).

#### In Vitro Cleavage with Furin

[^35^S]Met/Cys labeled pro-BMP was synthesized using rabbit reticulocyte lysate and proteins immunoprecipitated using antibodies specific for epitope tags in the protein. Digestion was carried out using recombinant furin (New England Biolabs) as described (19). All results were confirmed in a minimum of three independent cleavage assays.

#### Analysis of RNA

Total RNA was isolated from pooled ectodermal explants using TRIzol (Invitrogen). Semi qRT-PCR was performed as described (37, 38) using an annealing temperature of 56°C. qRT-PCR analysis of *HAMP* expression in Hep3B cells was performed as described (39)

#### Immunoblots of *Xenopus* extracts and Blastocoele fluid

Proteins were harvested from 10 pooled DMZs explants by freon extraction method as described previously (33). Fluid was aspirated and pooled from the blastocoele of the same number of gastrula stage embryos in each experimental group, as described(40). Proteins present in DMZs or in blastocoele fluid were denatured, deglycosylated with peptide-N-glycosidase F (PNGase F) as described (20). Proteins were resolved by SDS-PAGE under reducing and non-reducing conditions and transferred onto PVDF membranes. Membranes were probed with anti-actin (1:10000, Sigma), anti-pSmad1/5/8 (1:1000, Cell signaling) or anti-HA (3F10, 1:1000, Roche) antibodies. Immunoreactive proteins were detected using Enhanced Chemiluminescence reagent (Pierce) and light emissions captured with x-ray film. Images were scanned and relative band intensity was quantified using ImageJ software.

#### Transient Transfection and Immunoblot Analysis in Cultured Mammalian cells

HEK293T cells were purchased from American Type Tissue culture, authenticated at the University of Utah DNA sequencing core and tested for myco-plasma in the lab. Transient transfection and western blot analysis of HEK293T cells was performed as described (11) using anti-HA (3F10, 1:1000, Roche) and anti-actin (1:10000, Sigma) antibodies. Immunoreactive proteins were detected using Enhanced Chemiluminescence reagent (Pierce) and light emissions captured with x-ray film. Images were scanned and relative band intensity was quantified using ImageJ software. antibodies. 0.6M TMNK1 cells (purchased from the Japanese Collection of Research Bioresources Cell Bank (41)) were seeded to each well in 6-well plates and transfected with 1000ng of plasmid vector encoding BMP2, BMP2SIG, BMP2S2G, BMP6, BMP6R-G or empty vector using FuGENE® 4K Transfection Reagent (Promega). For co-transfection with BMP2 and BMP6, 500ng BMP2 and 500ng BMP6 plasmid vector were used for each well. Cells were cultured in transfection media (0.1 mL Opti-MEM and 0.9 mL DMEM with 10% serum per well) for one day, followed by growth media (1 mL DMEM with 10% serum per well) for the second day. Then, cells were cultured in 2 mL serum-free DMEM per well for 24 hours before collecting conditioned media. 0.8-1M Hep3B cells were seeded onto each well in 6-well plates and cultured with a mixture of 1mL conditioned media and 1mL MEM with 1% serum for 6 hours. Proteins from 1.8 mL conditioned media were precipitated by trichloroacetic acid and denatured in RIPA buffer for 5 minutes at 95°C under reducing and non-reducing conditions. TMNK1 and Hep3B cells were lysed by RIPA buffer with protease and phosphatase inhibitors. Protein densities of cell lysates were measured by colorimetric assay (Bio-Rad) and were denatured for 5 minutes at 95°C under reducing conditions. Proteins from cell lysates and conditioned media were separated by SDS-PAGE and then transferred onto nitrocellulose membranes. Membranes were probed with anti-pSMAD1/5/8 (1:1000 Abcam), anti-SMAD (1:1000 Abcam), anti-actin (1:5000 Sigma-Aldrich), or anti-HA-Tag (1:1000, Cell signaling). Immunoreactive proteins were visualized using the Syngene G: Box mini-imaging system.

#### Statistical analysis

Statistical analyses were performed using Student’s *t-*test or one-way analysis of variance (ANOVA) with Tukey or Holm-Sidak post-hoc test for multiple comparisons. The induction rates of *HAMP* and pSMAD in Hep3B cells were log transformed using natural log as the base before comparison. Differences with *P*<0.05 were considered statistically significant.

## Data Availability

All data described are contained within the manuscript.

## Supporting Information

This article contains supporting information

## Acknowledgements

*Funding and additional information*

This work was supported by the Eunice Kennedy Shriver National Institute of Child Health and Human Development, Grant/Award Numbers: 5R21HD102668 (JLC); National Institute of Diabetes and Digestive and Kidney Diseases, Grant/Award Number: R01DK128068 (JLC, JLB). This work utilized DNA and peptide shared resources supported by the Huntsman Cancer Foundation and the National Cancer Institute of the NIH (grant P30CA042014). The content is solely the responsibility of the authors and does not represent the official views of the NIH.

*Conflicts of interest*

There are no actual or perceived conflicts on the part of any author.

