## Supporting Figures 1-4 for "The BMP2 prodomain promotes dimerization and cleavage of BMP6 homodimers and BMP2/6 heterodimers in vivo"

Supporting Information included:

Figures S1-S4

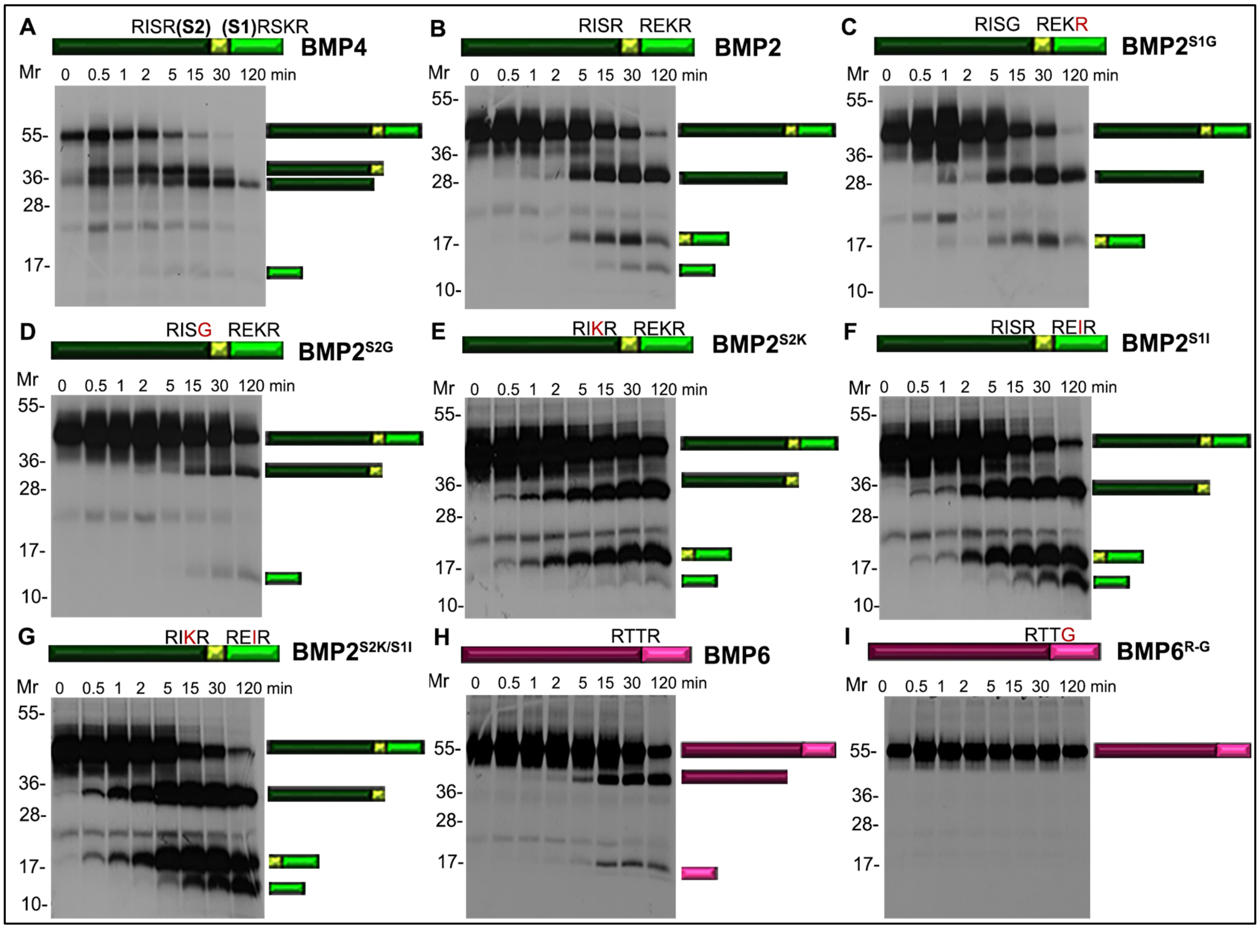


**Figure S1. In-vitro cleavage of wild type or mutant forms of BMP4, BMP2 and BMP6 precursor proteins.** (A-I) Radiolabeled precursor proteins were synthesized using rabbit reticulocyte lysate and incubated with recombinant furin. Aliquots were removed at different time points and analyzed by SDS-PAGE followed by autoradiography. Bands corresponding to the precursor, prodomain and ligand are indicated to the right of the gel. In vitro cleavage of BMP4 showing sequential cleavage at the S1 and then the S2 site (A). In-vitro cleavage of wild-type (B) and mutant forms of proBMP2 in which one of the two cleavage sites is mutated to prevent cleavage (C, D) or in which both sites are optimal (E; $BMP2^{S2K}$) or minimal (F; $BMP2^{S1I}$) furin motifs, or in which optimal and minimal furin motifs are switched (G; $BMP2^{S2K/S1I}$) (illustrated above each panel). In vitro cleavage of wild type (H) and cleavage mutant (I) forms of BMP6. Panels B-I are long exposures of the gels shown in Fig. 1A-H to show precursor, prodomain and cleaved ligand in a single panel. Gels shown in panels B-D, E-G or H-I were run on the same day and exposed for the same amount of time so that band intensities of cleavage products generated from wild type and mutant precursor proteins can be compared.

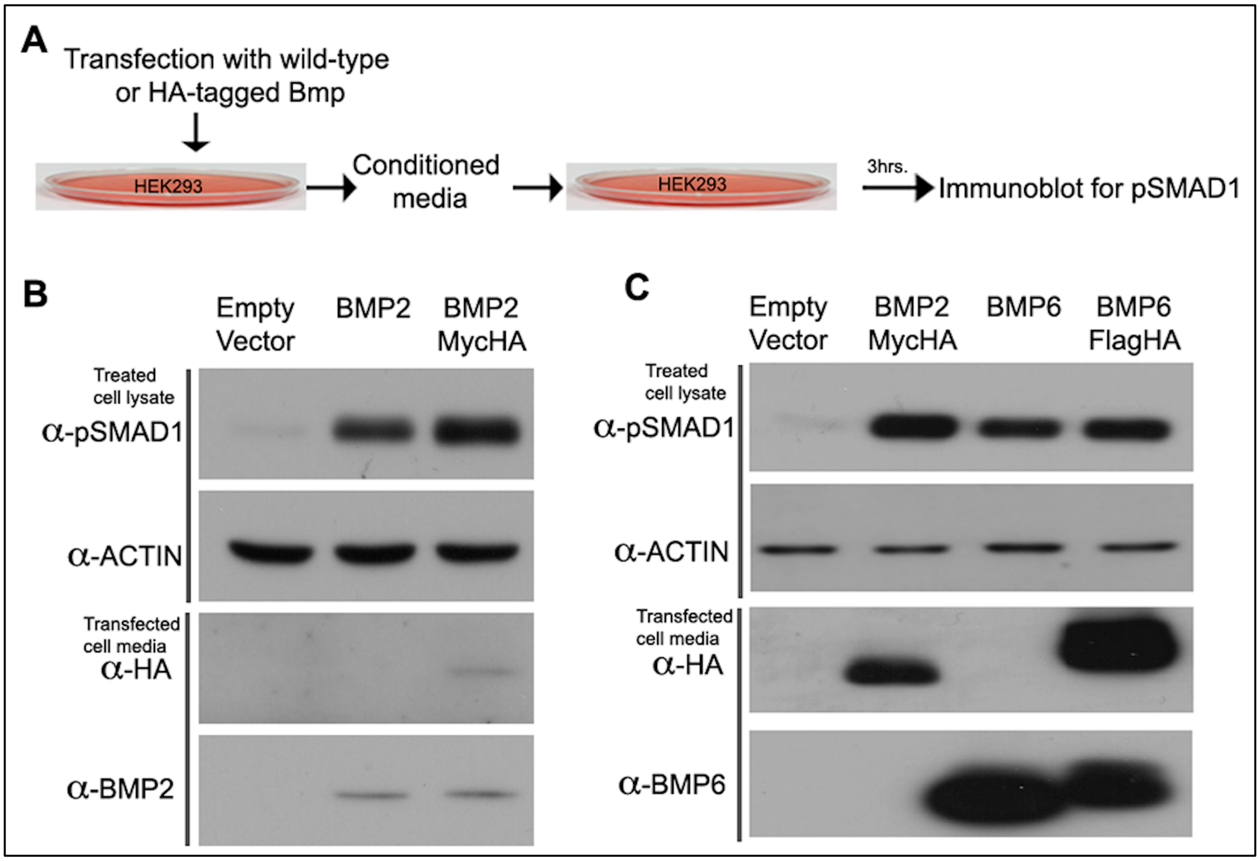


**Figure S2. Epitope tags in the ligand domain of BMP2 or BMP6 do not interfere with activity.** (A-C) HEK293 cells were transfected with cDNAs encoding empty vector, BMP2 or BMP6 with or without an HA tag in the ligand domain. Conditioned media was applied to untransfected cells for 3 hours after which cell lysates were collected and pSMAD1 levels analyzed by immunoblot. Immunoblots of cell lysates and conditioned media from transfected cells were probed with antibodies specific for the ligand domain or the HA tag in each BMP.

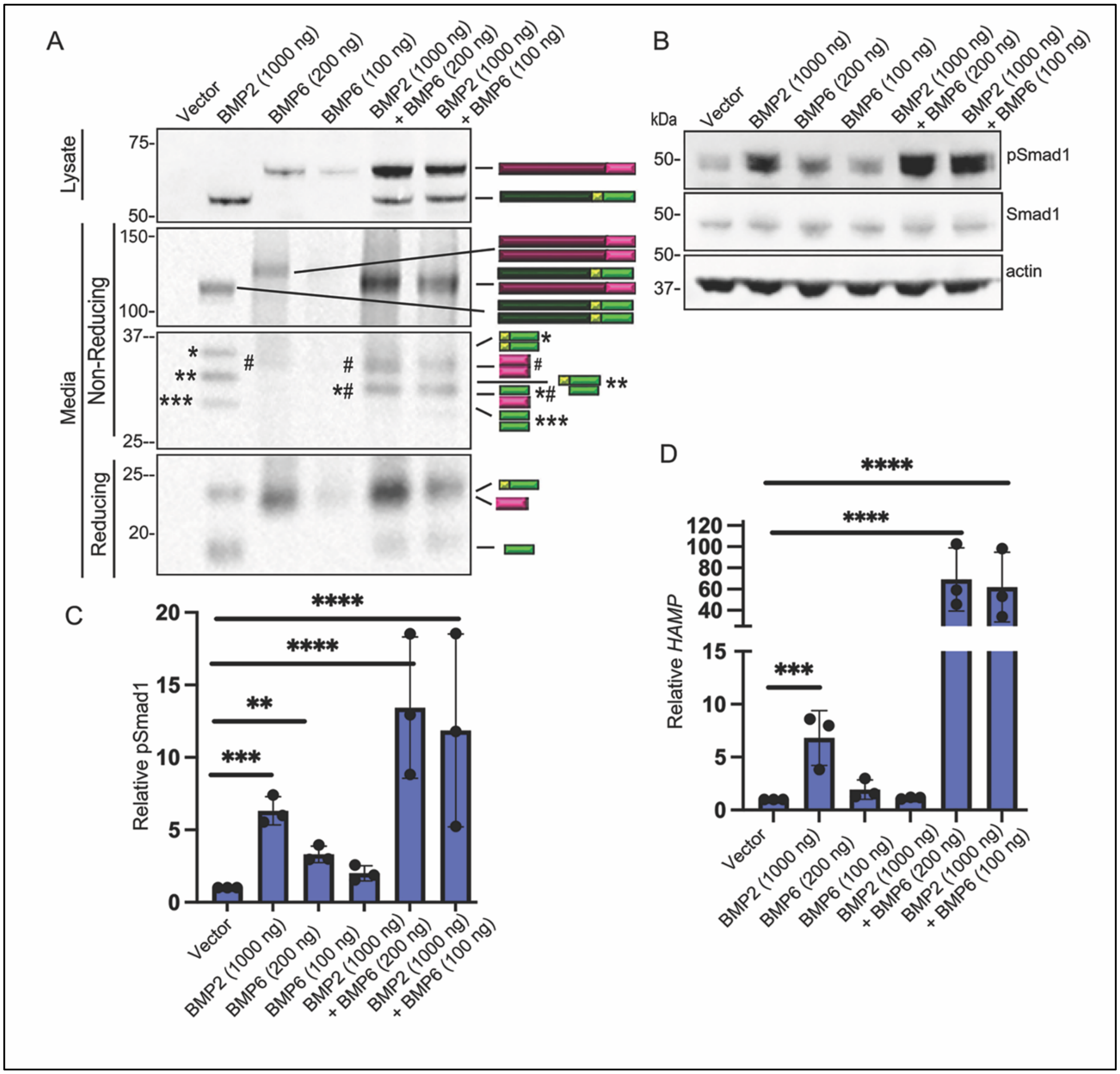


**Figure S3. Activity of BMP2 and BMP6 homodimers and heterodimers expressed in TMNK1 cells.** (A) TMNK1 cells were transfected with cDNAs encoding BMP2 or BMP6 precursors alone or together. Immunoblots of transfected TMNK1 cell lysates under reducing conditions and media under reducing or non-reducing (Non-Red) conditions probed with antibodies specific for the HA epitope in the ligand domain. Bands corresponding to precursor proteins and ligands illustrated to the right of the gel. (B-D) TMNK1 cells were transfected with cDNAs encoding BMP2 or BMP6 precursors alone or together. Conditioned media was collected 24 hours later and applied to

Hep3B cells for 6 hours, after which cell lysates and RNA were collected, pSMAD1 levels were analyzed by immunoblot, and *HAMP* levels were analyzed by qPCR. Representative pSMAD1 immunoblot (B), the relative level of pSMAD1 normalized to actin (C) or *HAMP* normalized to *RPL19* (D) and reported relative to that in cells transfected with empty vector are shown. Data are from a minimum of four independent experiments (mean ± SD) (**$p<0.01$, ***$p<0.001$, ****$p<0.0001$as determined by ANOVA with Tukey post-hoc test).

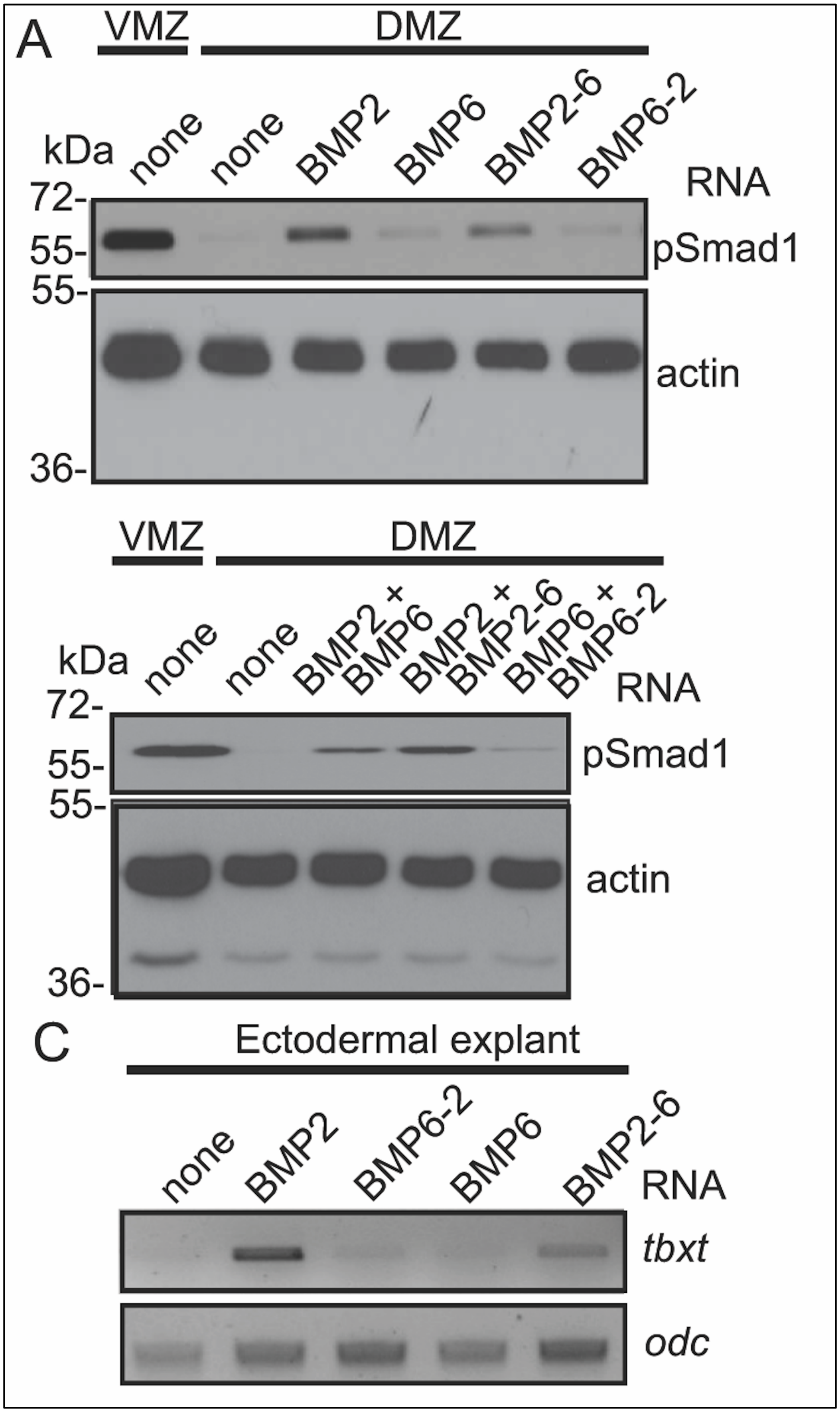

A
VMZ
DMZ
kDa
none
none
BMP2
BMP6
BMP2-6
BMP6-2
RNA
72-
55-
pSmad1
55-
actin
36-
VMZ
DMZ
kDa
none
none
BMP2 + BMP6
BMP2 + BMP2-6
BMP6 + BMP6-2
RNA
72-
55-
pSmad1
55-
actin
36-
C
Ectodermal explant
none
BMP2
BMP6-2
BMP6
BMP2-6
RNA
tbxt
odc

**Figure S4. The prodomain of BMP2 is necessary and sufficient to generate active BMP6 homodimers and BMP2/6 heterodimers in *Xenopus*.** (A, B) RNA encoding native or chimeric BMPs were injected individually (A) or together (B) near the dorsal midline of four-cell embryos. DMZ and VMZ explants were collected at stage 10 and pSmad1 levels were analyzed by immunoblot. Actin levels were analyzed in duplicate samples as a loading control. Data from these original blots was used to generate panels shown in Figure 3B and 4B. (C) BMP RNAs were injected alone or together into one animal pole blastomere of 2-cell embryos. Ectoderm was explanted at stage 11 and expression of *tbxt* was analyzed by semi qRT-PCR. Data from these original gels was used to generate panels on right shown in Figure 3E and 4E.
